# Sympathetic and parasympathetic involvement in time constrained sequential foraging

**DOI:** 10.1101/752493

**Authors:** Neil M. Dundon, Neil Garrett, Viktoriya Babenko, Matt Cieslak, Nathaniel D. Daw, Scott T. Grafton

## Abstract

Appraising sequential offers relative to an unknown future opportunity and a time cost requires an optimization policy that draws on a learned estimate of an environment’s richness. Converging evidence points to a learning asymmetry, whereby estimates of this richness update with a bias toward integrating positive information. We replicate this bias in a sequential foraging (prey selection) task and probe associated activation within two branches of the autonomic system, sympathetic and parasympathetic branches, using trial-by-trial measures of simultaneously recorded cardiac autonomic physiology. In general, lower value offers were accepted during periods of autonomic drive, both in the sympathetic (shorter pre-ejection period PEP) and parasympathetic (higher HF HRV) branches. In addition, we reveal a unique adaptive role for the sympathetic branch in learning. It was specifically associated with adaptation to a deteriorating environment: it correlated with both the rate of negative information integration in belief estimates and downward changes in moment-to-moment environmental richness, and was predictive of optimal performance on the task. The findings are consistent with a parallel processing framework whereby autonomic function serves both learning and executive demands of prey selection.

**Significance statement:** The value of choices (accepting a job) depends on context (richness of the current job market). Learning contexts, therefore, is crucial for optimal decision-making. Humans demonstrate a bias when learning contexts; we learn faster about improvements vs deteriorations. New techniques allow us to cleanly measure fast acting stress responses that might fluctuate with trial-by-trial learning. Using these new methods, we observe here that increased stress – specifically sympathetic (heart contractility) – might help overcome the learning bias (making us faster at learning contextual deterioration) and thereafter guide us toward better context appropriate decisions. For the first time we show that specific building blocks of good decision-making might benefit from short bursts of specific inputs of the stress system.

## Introduction

A specific but nonetheless ubiquitous value-based decision dilemma requires people to approach or avoid sequential offers that pit a reward against an opportunity cost of time, without full knowledge of what offers, if any, may follow. For example, by committing to a specific project, a contract worker receives payment (reward) while eschewing alternative projects during project completion (opportunity time cost), without knowing what alternative options will later emerge.

The Marginal Value Theorem (MVT; Charnov, 1976) proposes an optimality rule for such sequential decisions, whereby yields that exceed the average reward-rate (richness) of an environment should be approached and those falling below it should be avoided. The environment and its consequential opportunity cost thus prescribe choice selectivity; e.g., contract workers should only accept high yield projects (wiring a new supermarket) during construction booms when opportunity cost is high, and accept low yield projects (fixing a faulty domestic appliance) during construction downturns when opportunity cost is low. Accordingly, studies across various species (Cowie, 1977; McNamara and Houston, 1985), including humans (Hayden et al., 2011; Kolling et al., 2012), have predicted foraging behavior using MVT inspired models. Recent work in humans further resolves the computational challenge of learning dynamic environmental richness, by demonstrating that sequential choice behavior is best captured by an MVT inspired learning model. Specifically, both decisions to leave a patch and explore the environment in patch foraging (Constantino and Daw, 2015; Lenow et al., 2017) and capture behavior in prey selection (Garrett and Daw, 2019) adhere to the MVT predicted optimality policy that compares yields against fluctuating environmental richness which is learned via a standard delta rule. This later work further demonstrated that beliefs about environmental richness update with asymmetric bias, whereby improvements are learned at a higher rate than deteriorations; the ‘naïve perseverance of optimism’ (Garrett and Daw, 2019).

At present only one study has indirectly explored the relationship between stress related endocrine activity and foraging behavior by linking both acute and chronic stress elevation to overharvesting tendencies in patch foraging (Lenow et al., 2017). However, the main assays in that study (cortisol and self-report) probed stress fluctuations operating on longer time horizons than the faster acting learning needed to update beliefs about environmental richness. This time constant misalignment similarly affects a commonly used alternative assay of putative stress states - galvanic skin conductance - while further confounds such as arousal and spontaneous fluctuations complicate inferences regarding stress system contribution to pupillometry data (Bradley et al., 2008; Joshi et al., 2016; Krishnamurthy et al., 2017).

Measures of cardiac autonomic physiology have emerged as an exciting new approach for tracking rapid changes in cortically mediated stress responses fluctuating on a trial-by-trial basis but these have yet to be employed with sequential decision-making tasks. Such measures have nonetheless charted the effects of experimentally manipulated reward and difficulty on summary states of the sympathetic branch of the autonomic system, indexed with aggregated measures of beta-adrenergic myocardial mobilization (reviewed in Richter et al., 2016)). Of relevance to sequential decision-making, increased sympathetic states are associated with the difficulty of cognitive tasks and the relevance of reward – i.e., contractility increases with both increased difficulty and increased importance of reward (Richter et al., 2008; Kuipers et al., 2017). Further, where task difficulty is either unknown (Richter and Gendolla, 2009) or user-defined (Wright et al., 2002), (mirroring the situation in sequential decisions), sympathetic states uniquely track reward relevance, suggesting they may be involved with learning the opportunity cost of an environment. However, such a conclusion requires linking autonomic and environmental fluctuations over shorter time-scales.

Here, we employ state of the art cardiac analyses (Barbieri et al., 2005; Cieslak et al., 2018) on electrocardiogram (ECG) and impedance cardiogram (ICG) data recorded continuously while subjects performed a prey selection task, capturing trialwise modulation of sympathetic and parasympathetic contributions of the autonomic state. Using these trialwise indices, we address three questions; (1) how drive in the different stress systems aligns with choice policy and responds to changes in environmental richness; (2) how activation in these systems correlate with learning parameters; and (3) if activity in either autonomic system is associated with optimal task performance.

## Methods

### Participants

We recruited twenty subjects via word of mouth. Nine subjects were male and had a mean (standard deviation) age of 19.11 (1.37) while the remaining eleven female subjects had a mean (standard deviation) age of 20 (2.18). Total mean (standard deviation) age of our sample (N=20) was 19.65 (2.18). Two male subjects and one female subject reported themselves left-handed. All subjects provided informed consent to participate and experimental procedures were carried out following IRB approval from the University of California, Santa Barbara.

### Task

Subjects spent 24 minutes playing the Prey Selection task (Garrett and Daw, 2019); a computerized video game emulating a formal sequential foraging task under a time constraint (see Fig. 1A-C). Subjects were pilots of a space ship and instructed to harvest as much fuel as possible to earn a bonus ($0.01 per point). Fuel was harvested by capturing sequentially approaching space-invaders. Invaders carried either a high (80 points) or low (30 points) fuel reward, and a high (8s) or low (3s) capture cost. The four identities (see Fig. 1B) mapped onto a three-tier profitability rank: high (high reward / low cost); mid (high reward / high cost, or low reward / low cost); and low (low reward / high cost). Participants spent half of the game-time foraging in an environment with a disproportionately high concentration of high profit invaders (boom; high:mid:low = 4:2:1) and the other half in an environment with a disproportionately high concentration of low profit invaders (downturn; high:mid:low = 1:2:4) (see Fig. 1C). The two environments had different background colours, and subjects were informed that they would differ in terms of invader concentrations, but not explicitly how. Half of the subjects foraged in the order boom to downturn (BD) and the other half in the order downturn to boom (DB), with an opportunity to rest in-between the two environments.

**Figure 1:**
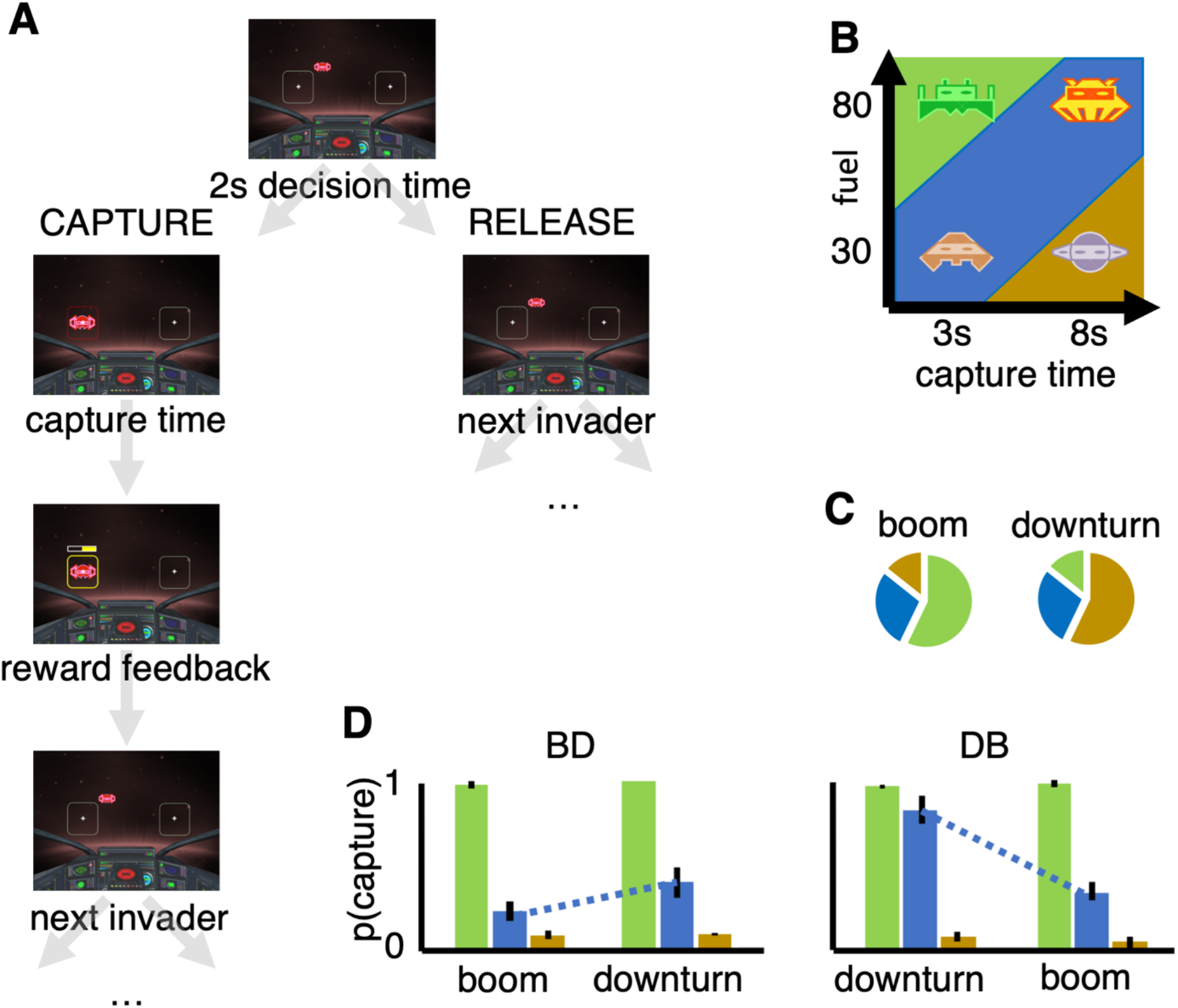
Prey selection paradigm. Subjects decide whether to capture or release serially approaching invaders during their two-second approach to the cockpit (panel A). Releasing an invader progresses immediately to the next invader, while capturing the invader incurs a capture time cost, and fuel reward. Four invader identities (panel B) map onto a two-by-two reward-bycost value space, and can be described categorically as hi (green), mid (blue), or low (brown) profitability. Subjects foraged for twelve minutes in each of two reward states states (panel C) with different proportions of invader profitability. Panel D: Replication of Garret and Daw (2019) learning asymmetry. Order of foraging (boom – downturn, BD; downturn – boom, DB) predicts optimal behavior (higher rank 3 captures in downturn relative to boom state). Learning deterioration of a reward state takes longer than learning state improvement. Error bars illustrate the standard error of the mean across subjects.

Subjects performed the Prey Selection task seated 150cm from a 68.6cm (diameter) computer monitor and registered responses on a standard PC keyboard. Experimental stimuli were presented on a Mac mini computer, using Psychtoolbox extensions (Brainard, 1997; Pelli, 1997; Kleiner et al., 2007) in MATLAB v9.4 (MATLAB, 2018a).

The task paradigm is described in Figure 1A. At all times subjects saw a cockpit with two target boxes, one on the left and one on the right of the screen. On each trial, subjects had two seconds to decide if they wished to capture or release an invader approaching their cockpit. Invaders would pseudo randomly approach one of the two target boxes. Captures were registered by holding the response button corresponding to the target box in the path of the incoming invader (z with left index finger for the box on the left, m with right index finger for the box on the right). Subjects were required to keep the response button held as the invader finished its two second approach to the box, and thereafter for the entirety of the capture time (2s or 7s). Following a successful capture, a feedback screen (1s) described the harvested reward, during which subjects could release the response button. After the feedback screen the next invader immediately began its approach. Releases were registered by holding the response button corresponding to the response box opposite the incoming invader. Subjects were required to hold the response button until the invader reached the capture box. The invader would then disappear and the next invader would begin its approach. Errors carried an 8s time penalty, during which the response boxes disappeared and no invaders approached. Errors were (i) failing to register any response during the 2s invader approach; (ii) releasing the response button before the invader reached the response box (captures and avoids); (iii) releasing the response button before the end of the capture cost (captures only). Note that requiring the full approach time on both captures and releases, regardless of the latency of response execution, subjects can only use choice policy and not vigor (see Guitar-Masip et al., 2011) to optimize performance. Also, trialwise pseudorandom mapping of capture and release onto either hand reduced the confounding influence of action hysteresis on decisions (see Valyear et al., 2019). Finally, to encourage full exploration of each environment, 25% of trials were forced-choice. On these trials, a red asterisk would appear above one of the two capture boxes, and participants were instructed to press this response button regardless of whether they wished to perform the corresponding capture or release. Error and forced-choice trials were excluded from later analyses. Participants received a standardized set of instructions, performed a two-minute block of practice trials and required a perfect score on a questionnaire probing task comprehension before starting the task.

### Physiological recording and preprocessing

Physiological measures of ICG and ECG were collected using non-invasive approaches with a total of ten EL500 electrodes. Prior to each electrode placement, an exfoliation procedure was performed on each electrode location to maximise signal quality. An approximate one-inch area of skin was cleaned with an abrasive pad, followed by exfoliation with NuPrep gel (ELPREP, BIOPAC). Once the skin area was fanned dry, a small amount of BIOPAC GEL100 was placed on the electrode and on the skin. In order to assess ICG, a total of eight of the ten electrodes were placed on the neck and torso: two on each side of the neck and two on each side of the torso (Bernstein, 1986). ECG recordings were obtained with a total of two sensors placed beneath the right collarbone and just below the left ribcage. The ICG electrodes provided the necessary ground. Continuous ECG was collected using an ECG100C amplifier and continuous ICG using a NICO100C amplifier (both from BIOPAC). Data were integrated using an MP150, system (BIOPAC) and displayed and stored using AcqKnowledge software version 4.3 (BIOPAC). Both ECG and ICQ timeseries were recorded at 1000Hz. We recorded raw ECG (ECG) and both the raw (z) and derivative 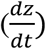 of the ICQ; the latter facilitates the identification of key impedance inflection points required to estimate the pre-ejection period (PEP). Both z and 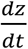 were high pass filtered to remove respiratory artefact. Below, reference to continuous ICG refers to 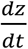.

We extracted three estimates of the physiological state at each heartbeat (see Fig. 2) – pre-ejection period (PEP), high frequency heart rate variability (HF HRV) and heart rate (HR). Semi-automated software MEAP labeled the continuous ECG and ICG (Cieslak et al., 2018). For each heartbeat, the ECG R point serves as the t=0 landmark for within-heartbeat events. The time interval between the ECG Q point and the ICG B point (see Fig. 2) defines the pre-ejection period (PEP) of a heartbeat, which is related to the contractility of the heart muscle before blood is ejected. However, due to difficulty in reliably capturing the relatively small Q point, PEP is often calculated as the difference between the easily detected ECG R point and the ICG B point (the RBI). This latter interval is comparable to PEP in reliability (Kelsey et al., 1998, 2007) and validity (Kelsey et al., 1998; Mezzacappa et al., 1999) and sometimes referred to as PEPr (Berntson et al., 2004). We used the RBI definition for our measure of PEP. Reduced values of PEP reflect a shorter pre-ejection interval, indicating increased sympathetic cardiovascular drive. However, to align PEP with the direction of our other autonomic variables, allowing easier comparison from the results of later analyses, we negative signed (i.e., *-1) all extracted values. For all subsequent references to PEP, higher values reflect increased sympathetic cardiovascular drive.

**Figure 2:**
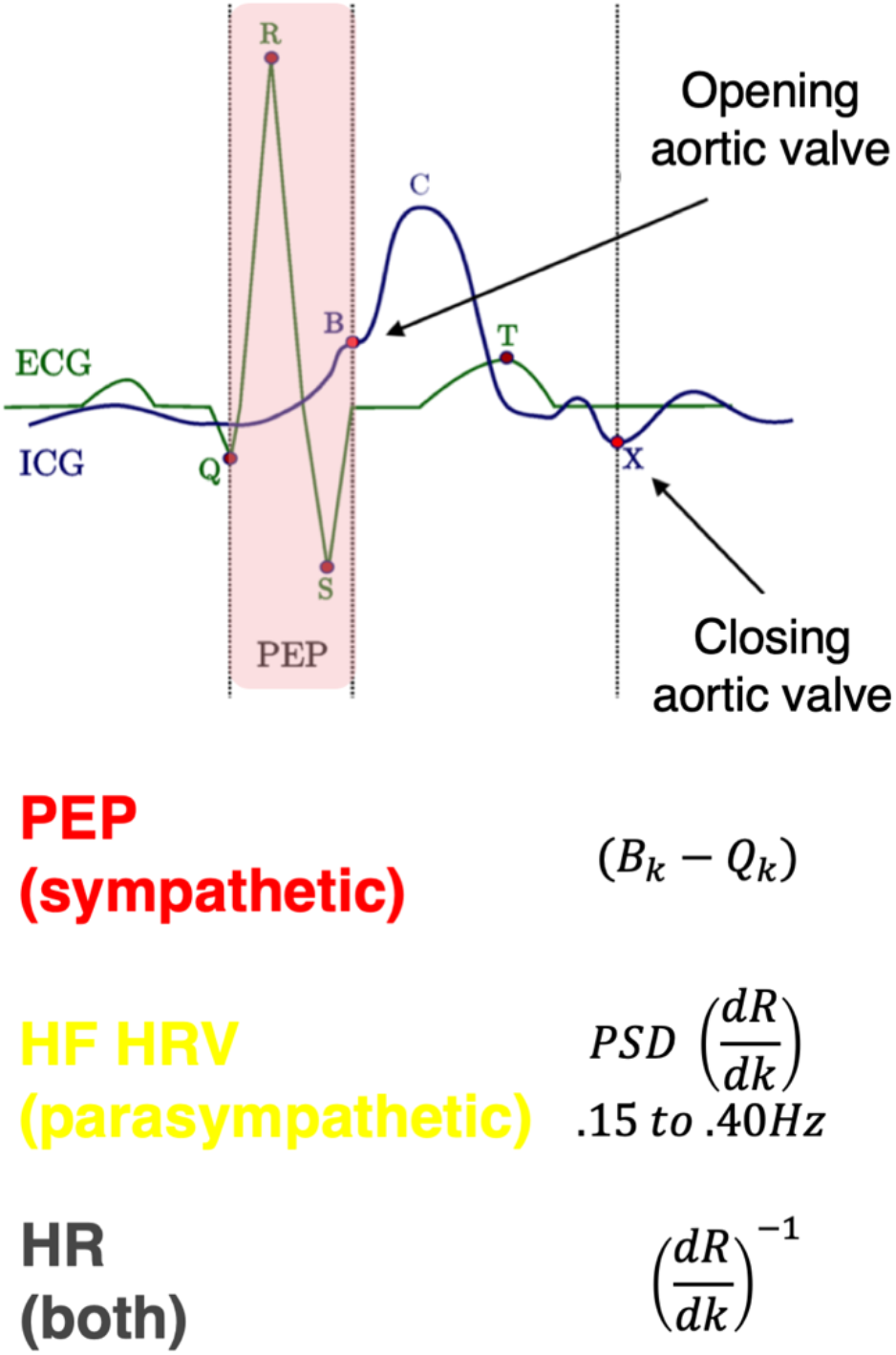
Dynamics of a template heart beat (k), as measured by electrocardiogram (ECG; green) and impedance cardiogram (ICG; blue). Pre-ejection period (PEP) indexes sympathetic mediated myocardial contractility, computed as the time between early ventricular depolarization (point Q on the ECG) and the opening of the aortic valve (point B on the ICG). Note that in our analyses we use a more easily identified ECG landmark for early ventricular depolarisation (point R) and reverse signed each estimate (see Methods: Physiological recording and preprocessing). High frequency heart rate variability (HF HRV) assays vagal (parasympathetic) input, computed as the power spectrum density (PSD) of R-R intervals, between 0.15 and 0.40Hz. Heart rate (influenced by both sympathetic and parasympathetic activity) is computed as the reciprocal of the R-R intervals.

Variability in R-R intervals are commonly separated into low (0.04 – 0.15Hz) and high (0.15 – 0.40Hz) frequencies, with the former reflecting a complex mixture of sympathetic and parasympathetic tone (Saul et al., 1991) and the latter serving as a proxy of parasympathetic cardiac tone (Barbieri et al., 2005). We characterized high frequency heart rate variability (HF HRV) with point process estimation software developed by Riccardo Barbieri and Luca Citi, executed as MATLAB code (available at http://users.neurostat.mit.edu/barbieri/pphrv). In brief, the code uses inverse Gaussian regression and spectral analysis of the latencies of ECG R points to generate a continuous estimation of HF HRV (Chen et al., 2007), i.e., power spectral density between 0.15 and 0.40 Hz, expressed in *ms*^2^/*Hz*. We extracted the logarithmic transform of these pointwise estimations as a proxy of parasympathetic input to the heart, with higher values reflecting increases in this autonomic branch.

Finally, we used the reciprocal of R-R intervals as a measure of heart rate (HR), such that higher HR values reflect a decrease in the interval between R points, i.e., increased heart rate. While heart rate is influenced by both sympathetic and parasympathetic inputs, we included it in our analyses to control for a known cardiac effect where increased left-ventricular preload time (which occurs with slowing HR) can shorten PEP independent of sympathetic influences (Sherwood et al., 1990).

Trialwise estimates of the three physiological states were next derived by taking an average from all heartbeats during the two-second time window while invaders approached the spaceship on each trial. This provided trialwise estimates of the physiological states during a uniform length time window for all trials, regardless of the executed decision or the identity of the invader.

Finally, each trialwise physiology estimate was corrected for trialwise respiratory state. We performed this additional trialwise respiration correction to account for known influences of respiratory activity on heart rate and vagal tone (Larsen et al., 2010), computed in our pipeline from raw ECG R points. We defined trialwise respiratory state as the average normalized product of the phase and magnitude of respiration activity (estimated from the z time series) at each R point during the two-second time window of invader approaches. Respiration corrected measures of each trialwise physiological state were the residuals from a linear model of each raw trialwise physiology state and the trialwise respiratory state, performed separately for each subject.

We will now present the statistical analysis methods and results separately for three separate branches of analyses. The first branch uses ANOVA and linear mixed effects models to explore the dynamics between physiological states, choices and objectively estimated measures of environmental richness and its moment-to-moment derivatives. The second analysis employs computationally modelled subjective estimates of the environment’s richness; and compares models that allow the learning parameter to vary with physiological states. In the third and final analysis we test if blockwise changes in physiological state predict optimal task performance.

### Analysis branch 1 - Methods

All ANOVAs and trialwise mixed-effects models were fitted using lme4 (Bates et al., 2015) and lmerTest (Kuznetsova et al., 2017) packages in R. Unless otherwise specified, trialwise logistic models were fitted with logit link functions and Laplacian maximum likelihood approximation, while trialwise models of continuous measures used restricted maximum likelihood approximation (REML). Also, unless otherwise specified, each trialwise mixed-effects model fitted a fixed effect for each specified coefficient, and an individual intercept for each subject. For both ANOVA and trialwise mixed-effects models, significant marginal effects of significant higher order interactions are not reported. Post-hoc ANOVA contrasts use Tukey correction. All other data pre-processing and analyses were conducted using MATLAB v9.4 (MATLAB, 2018a).

### Analysis branch 1 - Results

Our first behavioral analysis attempted to replicate the Garrett and Daw (2019) finding regarding asymmetric belief updating. To this end, we ran a three-way mixed ANOVA of mean capture-rate as a function of between-group factor *order* (RP, PR) and two repeated-measures factors *env* (boom, downturn) and *rank* (hi, intermediate, low). Summarized in Figure 1D, the ANOVA reported a significant three-way interaction between *order, env* and *rank* (*F* = 3.49, *df* = (2,90), *p* = 0.035). We accordingly contrasted mean capture rates between BD and DB, for the six levels of the *env*rank* interaction. These contrasts demonstrated significantly higher capture of mid-rank invaders in the downturn environment for order DB, relative to BD (*μ* = 0.828 vs. *μ* = 0.402, both *s. e*. = 0.050, *p* < 0.001), with no other contrasts reaching statistical significance. In other words, foraging during the two order conditions differed only in terms of adjustments in the number of mid-rank captures in the downturn environment, in line with Garrett and Daw (2019), and suggestive of slower learning in the face of a contextual deterioration.

A corollary of asymmetric belief updating (prioritizing positive information) is a decision threshold weighted preferentially toward recent reward over recent cost. To formally probe the influence of current and recent offers on choice behavior, we fitted a trialwise mixed-effects model of *choice_t_* (capture, release) as a function of an intercept term and four parameters: reward and delay on a given trial *t* (respectively: *reward_t_, delay_t_*) and reward and delay on the previous trial, i.e., *t* − 1 (respectively: *reward_t−1_, delay_t−1_*). The model yielded a significant positive influence of *reward_n_* (*β* = 2.20, *s. e*. = 0.071, *p* < 0.001) and significant negative influence of *delay_n_* (*β* = −2.19, *s. e.*= 0.073, *p* < 0.001) on capture probability. Regarding recent offers, choice was only influenced by recent reward; *reward_t−1_*: (*β* = –0.340, *s. e.* = 0.067, *p* < 0.001), *delay*_*t*−1_: (*β* = 0.062, *s.e.* = 0.067, *p* = 0.353). The negative coefficient for *reward_t−1_* is predicted by MVT; for example a high reward drives positive belief updating (of the environment’s richness), increasing opportunity cost to future captures, and decreasing future acceptance. The specific influence of *reward_t−1_* further supports the Garrett and Daw (2019) finding that positive information integrates more readily into state appropriate behavioral policy, relative to negative information.

We next assessed the influence of current physiological state on the relationship between current offer and choice behavior. In three separate models, i.e., one each for PEP, HF HRV and HR, we ran a trialwise mixed-effects model of *choice_t_* (capture, release) as a function of the two-way interaction effects *phsyiology_t_*reward_t_* and *physiology_t_*delay_t_*. We also included, in each model, an intercept term, and nuisance coefficients *order* (BD, DB), *env* (boom, downturn) and *trial index*. In line with the behavior analyses above, each model showed the same influence of *order* and *state* on choice – higher capture rates in the downturn state, and for the DB order (all p-values < 0.004). Each model also returned a significant negative coefficient for *trial index* (all p-values < 0.001) reflecting higher capture rates early in foraging.

In addition, the PEP model returned a significant positive coefficient for the *physiology*delay* effect (*β* = 0.548,5. *e*. = 0.147, *p* < 0.001), suggesting that an increase in contractility (shorter PEP) blunts the negative association between delay and capture. The *physiology*reward* did not reach statistical significance (*β* = −0.241, *s. e.* = 0.148, *p* = 0.098).

The HF HRV model also returned a significant positive coefficient for the *physiology*delay* effect (*β* = 1.09, *s. e.* = 0.145, *p* < 0.001), suggesting that the negative association between delay and capture may be alleviated with increasing HF HRV power (increased parasympathetic tone). In contrast to PEP, the *physiology*reward* effect from the HF HRV model was also significant (*β* = −1.29, *s.e*. = 0.144, *p* < 0.001); the negative coefficient suggests the positive association between reward and capture also alleviates when HF HRV power increases.

The HR model returned a significant negative coefficient for the *physiology*delay* effect (*β* = −0.892, *s.e.* = 0.168, *p* < 0.001), and a significantly positive coefficient for the *physiology*reward* effect (*β* = 1.14, *s. e.* = 0.169, *p* < 0.001), suggesting that decreased heart rate blunts both the aversion of delay and the appeal of reward.

The findings from these preliminary models suggest that increases in both branches of the autonomic state (PEP and HF HRV) are aligned with lower value acceptance, with PEP tuned more specifically to the cost dimension of value. HR, in contrast, decelerates during moments of low value capture. A shortcoming of these models, however, is that they do not consider value relative to the current rate of reward, i.e., environmental richness. Also, by running three separate models, i.e., one each for the three different physiology variables, we cannot confirm if physiological associations with reward and cost are independent of one another. We accordingly simplified the parameter space such that a single trialwise parameter *value_t_* would account for both dimensions of value, adjusted by an evolving estimation of the opportunity cost at the time of each choice.

*value* for a given trial *t* was defined as:

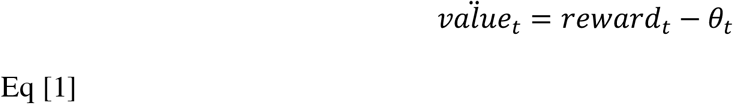

Where *reward_t_* is the reward of the offer on trial *t* and *θ_t_* is the opportunity cost of capturing offer *t*, computed as:

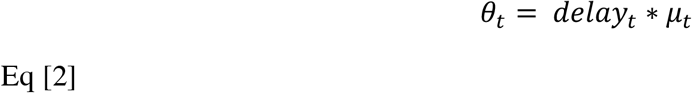

Where *μ_t_* is the average rate of reward captured per second through to the beginning of trial *t*, i.e.:

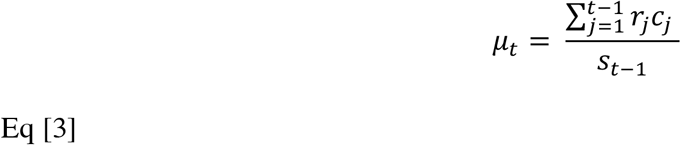

where *c_j_* is a discrete variable reflecting the selection for trial *j* (1 = capture, 0 = release) and *s_t−1_* is the time in seconds at the end of the feedback of trial *t* – 1 (relative to opening frame of the first trial of the experiment).

Accordingly, *value* approaching positive ∞ reflect offers that exceed the opportunity cost of the current moment, and should readily be captured, while *value* approaching negative ∞ describe readily avoidable offers where the reward harvested in exchange for the absorbed time cost is not justified at that moment. In choice models, this relationship is reflected by a positive coefficient, i.e., higher choice probability follows higher *value.*

We then ran a single model of *choice_t_* (capture, release) as a function of three two-way interaction effects: *PEP_t_*value_t_*, *HF_t_*value_t_* and *HR_t_*value_t_*, in addition to an intercept term, and nuisance coefficients *order* (BD, DP), *env* (boom, downturn) and *trial index.* The model (see Fig. 3A) returned a significantly negative coefficient for the *PEP* * *value* interaction (*β* = – 0.306,5. *e.* = 0.134, *p* = 0.023), suggesting that increased contractility blunts the positive association between choice and *value;* in other words, sympathetic drive is associated with higher acceptance of offers presenting low value at a given moment. The model also returned a significantly negative coefficient for the *HF * value* interaction (*β* = –0.861, *s. e.* = 0.130, *p* < 0.001), which also describes a blunting of the *value* coefficient when HF HRV power increased; in other words, parasympathetic drive is also associated with higher acceptance of low value offers at a given moment. The *HR*value* interaction also reached statistical significance (*β* = 0.341, *s.e*. = 0. 144, *p* = 0.018), indicating that a slower heart-rate again blunts the positive association between value and capture.

**Figure 3:**
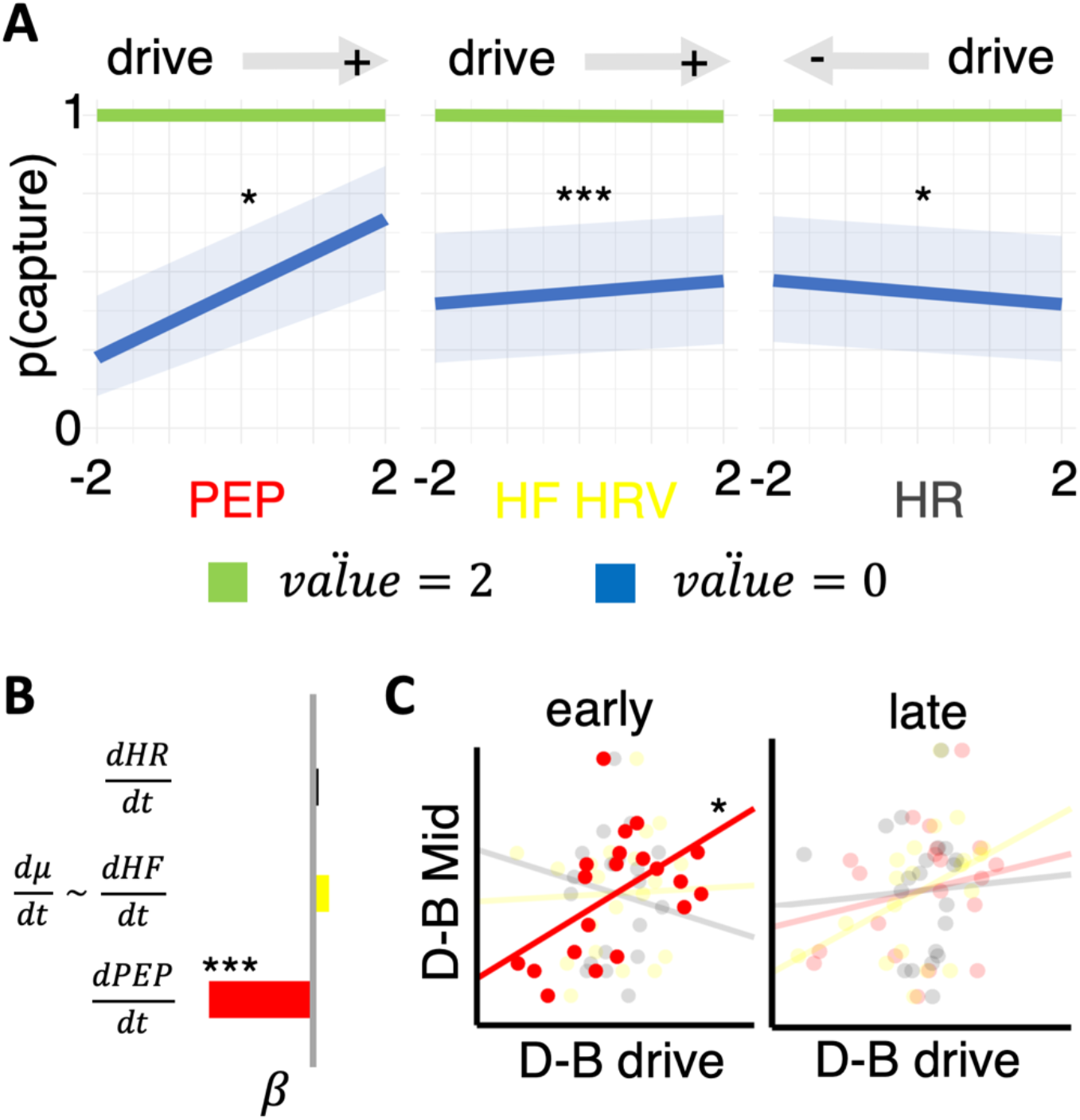
Relationship between autonomic states, value and capture. Panel A: From a single model (see analysis branch 2) a series of significant two-way interactions is described where increased contractility (shorter PEP) and increased vagal tone (higher HF HRV) are both associated with increased capture of lower value (blue line) offers. In contrast, increased low value capture is associated with decreased heart-rate (HR). Here, value is described as in equation 1, i.e., reward relative to opportunity cost in current objective reward state, with two levels of value (0 = low; 2 = high) selected from this continuous variable for illustration. As a guide, grey arrows above the plot describe whether increased capture for low value is associated with increased (+) or decreased (−) drive within the respective autonomic state. Panel B: PEP tracks deterioration in environmental richness. Only trialwise changes in PEP 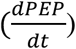 significantly predicted trialwise changes in environmental richness 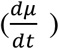, positive coefficient indicates that decreases in reward rate, i.e., deterioration, increased contractility (shorter PEP). Panel C: PEP predicts optimal learning. Blockwise changes in percentage of rank 3 captures (downturn – boom) modeled as a function of blockwise changes in PEP (red), HF HRV (yellow) and HR (grey); separately for mean physiological changes in the early (0 – 360s) and later (360-720s) portion of blocks. Optimal performance (higher D-B rank 3 score) predicted by higher relative drive in early downturn state, relative to early boom, only for PEP. \**p*<0.05; \*\*\**p*<0.001

Sympathetic and parasympathetic drive may track with unpleasant decisions, i.e., a specific action associated with an individual offer and its immediate negative consequence. However, mobilization of either of these autonomic branches could also reflect their association with derivatives (perturbations) of rate of reward, i.e., learning the richness of the environment. We consider *μ_t_* from Eq [3] a proxy for the evolution of this reward rate. The derivative of *μ_t_* with respect to trial *t*, i.e.,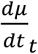, represents trialwise perturbations. We modelled 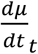 as a function of 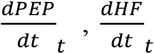, and 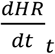. The model also contained an intercept term and nuisance coefficients *env, choice, order*, and *trial index*.

This model of 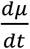 (see Fig. 3B) returned a significantly negative coefficient for 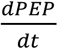 (*β* = –0.006, *s. e.* = 0.002, *p* < 0.001), indicating countercyclical perturbations in contractility and reward rate – i.e., if the reward rate decreased, contractility would increase, and vice versa. The coefficients for *HF* and *HR* failed to reach statistical significance (both p-values > 0.591). Perturbations in the reward rate are therefore exclusively associated with sympathetic drive; contractility increases as reward rate reduces. A control model of rectified reward rate derivative, i.e., 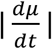, modelled as a function of 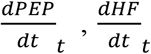 and 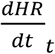 returned no significant physiology coefficients (all p-values > 0.079), ruling out a more global responsivity of the sympathetic system to reward rate changes in any direction.

Finally, we reran all models in Analysis branch 1 using the raw physiology measures, i.e., not corrected for respiration state using the residualization procedure outlined in the pre-processing section. The pattern of results were the same across all models.

### Analysis branch 2 - Methods

The key finding from our analysis so far centers on perturbations in the sympathetic state tracking perturbations in the reward rate. This may reflect a relationship between the sympathetic state and learning, however the objective measure of reward rate used above (the rate of reward harvested per second) is not direct evidence that subjects are learning these perturbations in environment quality. In this branch of analysis, we employ computational models fitted to the choice data, and estimate parameters of the subjective estimate of reward rate, in addition to parameters scaling the learning of perturbations in the reward rate. Recall from the previous section, that *μ_t_* represents a moving threshold against which encountered options can be assessed. A simple means by which participants can keep track of *μ_t_* (McNamara and Houston, 1985; Hutchinson et al., 2008; Constantino and Daw, 2015) is to implement an incremental error-driven learning rule (Schultz et al., 1997; Sutton and Barto, 1998) whereby a subjective estimate of *μ_t_*, which we will refer to here as *ρ* for clarity, is incrementally updated according to recent experience. The optimal policy from the MVT remains the same: capture an option *i*, whenever the reward, *r_i_*, exceeds the opportunity cost of the time taken to pursue the option. As with Eq [2], opportunity cost is calculated as the time, *delay_i_*, that the option takes to pursue (in seconds) multiplied by the subjective estimated of the reward rate, *ρ*.

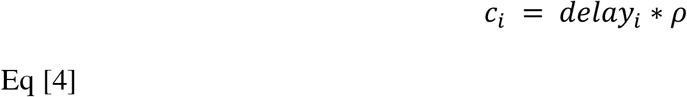

We will refer to this measure of opportunity cost, using the subjective estimate *ρ*, as *c_i_*. As with the objective measures, participants should capture items that exceed their subjective estimate of the reward rate, i.e., whenever *r_i_* ≥ *delay_i_* * *ρ*. Note that we assume quantities *r_i_* and *delay_i_* are known to participants from the outset since they were easily observable and each of the 4 invader identities (*i* = {1,2,3,4}) always provided the exact same *r_i_* and *delay_i_*.

We assumed that subjects learn *ρ* in units of reward, using a Rescorla-Wagner learning rule (Rescorla and Wagner, 1972; Sutton and Barto, 1998) which is applied at every second (*s*). After each second, their estimate of the reward rate updates according to the following rule:

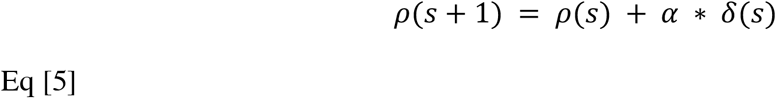

Here, *δ*(*s*) is a prediction error, calculated as:

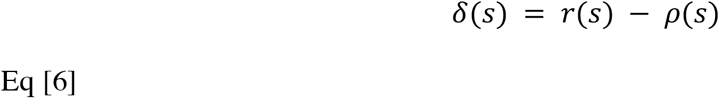

where *r*(*s*) is the reward obtained. *r*(*s*) will either be 0 (for every second in which no reward is obtained, i.e. during approach time, capture time and timeouts from missed responses) or equal to *r_i_* (following receipt of the reward from captured option *i*).

The learning rate *α* acts as a scaling parameter and governs how much participants change their estimate of the reward rate (*ρ*) from one second to the next. Accordingly, *ρ* increases when *r*(*s*) is positive (i.e. when a reward is obtained) and decreases every second that elapses without a reward.

### Symmetric and Asymmetric Models

We first implemented two versions of this reinforcement learning model, used previously to test for the presence of learning asymmetries in this task (Garrett and Daw, 2019). A *Symmetric Model*, with only a single *α* and a modified version, an *Asymmetric Model*, which had two *α: α*^+^ and *ρ*^−^. In this second model, updates to *ρ* apply as follows:

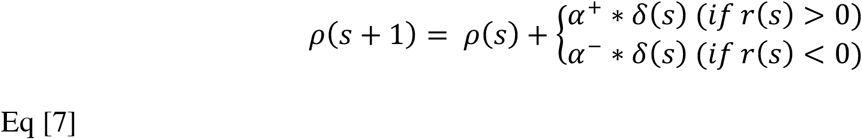

This second model allows updates to occur differently according to whether a reward is received or not. We refer to the mean difference in learning rates as the *learning bias* (*α*^+^ − *α*^−^). A positive learning bias (*α*^+^ > *α*^−^) indicates that participants adjust their estimates of the reward rate to a greater extent when a reward is obtained compared to when rewards are absent. The converse is true when the learning bias is negative (*α*^+^ < *α*^−^). If there is no learning bias (*α*^+^ = α^−^) then this model is equivalent to the simpler Symmetric Model with a single *α*.

### Asymmetric Models with physiology

Next, we extended the Asymmetry Model described above to test whether learning from positive (*α*^+^) or negative (*α*^−^) information was modulated by trial to trial perturbations in the physiological state. For each physiological state (i.e., HR, PEP and HF HRV) we compared two separate models (i.e., six models were fitted in total).

For each physiological state, the first model (Asymmetry Phys *α*^+^) tested physiological modulation of the *α*^+^ parameter. In this model, *α*^+^ was adjusted at each moment in time, according to the participants physiological state on that trial:

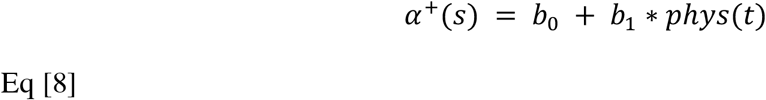

Here, *t* indexes the current trial. *b*_1_ governs the extent to which the learning rate for positive information (*α*^+^) is adjusted by the trialwise measure of the physiological state. *α*^−^ was unmodulated by physiological state in this model.

The second of these additional models (Asymmetry Phys *α*^−^) tested physiological modulation of the *α*^−^ parameter. This was setup exactly as for Asymmetry Phys *α*^+^, except *α*^−^ was adjusted on each trial, according to:

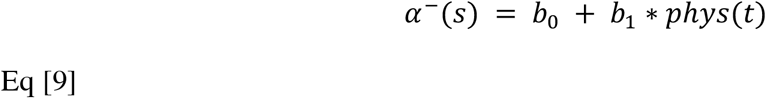

Here, *b*_1_ governs the extent to which the learning rate for negative information (*α*^−^) is adjusted on each trial. *α*^+^ was unmodulated by physiological state in this model.

In all models, the probability of capturing an item is estimated using a softmax choice rule, implemented at the final frame of the encounter screen as follows:

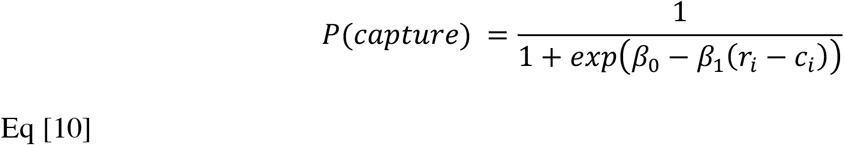

This formulation frames the decision to accept an option as a stochastic decision rule in which participants (noisily) choose between two actions (capture/release) according to value of each action. The temperature parameter *β*_1_ governs participants’ sensitivity to the difference between these two values whilst the bias term *β*_0_ captures a participant’s general tendency toward capture/release options (independent of the values of each action). Note that under the above formulation, negative values for *β*_0_ indicate a bias towards capturing options, positive values indicate a bias towards releasing options.

In each model, *ρ* was initialized at the beginning of the experiment to the arithmetic average reward rate across the experiment, but subsequently carried over between environments (Constantino and Daw, 2015; Garrett and Daw, 2019). For each participant, we estimated the free parameters of the model by maximizing the likelihood of their sequence of choices, jointly with group-level distributions over the entire population using an Expectation Maximization (EM) procedure (Huys et al., 2011) implemented in the Julia language (Bezanson et al., 2012) version 0.7.0. Models were compared by first computing unbiased marginal likelihoods for each subject via subject-level cross validation and then comparing these likelihoods between models (Asymmetric versus Symmetric) using paired sample ttests.

To formally test for differences in learning rates (*α*^+^, *α*^−^) we estimated the covariance matrix 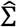 over the group level parameters using the Hessian of the model likelihood (Oakes, 1999) and then used a contrast 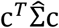 to compute the standard error on the difference *α*^+^ – *α*^−^.

### Analysis branch 2 – Results

A key feature of the learning previously observed in this task (Garrett and Daw, 2019) is that individuals adjusted their subjective estimates of the reward rate to a greater degree when the update was in a positive compared to negative direction. This *learning asymmetry* accounted for the order effect whereby participants changed capture rates between environments to a greater degree when richness improved (participants transitioned from downturn to boom) compared to when richness deteriorated (participants transitioned from boom to downturn), an effect we also observe here (see Fig. 1D).

First, we tested if the same learning asymmetry was present in our data by fitting choices to two reinforcement learning models (Sutton and Barto, 1998): a *Symmetric Model* and an *Asymmetric Model,* exactly as done previously (Garrett and Daw, 2019). Both models used a delta-rule running average (Rescorla and Wagner, 1972) to update *ρ* according to positive and negative prediction errors. Negative prediction errors were generated every second that elapsed without a reward (for example, each second of a time delay). Positive prediction errors were generated on seconds immediately following a time delay when rewards were received. The difference between the Symmetric Model and the Asymmetric Model was whether there were one or two learning rate parameters. The Symmetric Model contained just a single learning parameter, *α*. This meant that *ρ* updated at the same rate regardless of whether the update was in a positive or a negative direction. The Asymmetric Model had two learning parameters: *α*^+^ and *α*^−^. This enabled *ρ* to update at a different rate, according to whether the update was in a positive (*α*^+^) or a negative (*α*^−^) direction.

Replicating past findings (Garrett and Daw, 2019), the Asymmetric Model again provided a superior fit (see Table 1) to the choice data than the Symmetric Model (*t*(19) = 3.14, *p* = 0.005, paired sample ttests comparing Leave One Out cross validation scores for the Asymmetry versus the Symmetric Model) with information integration again being biased in a positive direction (*α*^+^ > *α*^−^. *z* = 1.80, *p* < 0.05 one-tailed). Prediction errors that caused *ρ* to shift upwards (following receipt of a reward) had a greater impact than prediction errors that caused *ρ* to shift downwards (following the absence of a reward).

**Table 1:** behavioural model fitting and parameters. model fitting and parameters for Symmetry and Asymmetry Models. The table summarizes for each model its fitting performances and its average parameters: LOOCV: leave one out cross validation scores, summed over participants; α: learning rate for both positive and negative prediction errors (Symmetric Model); α+: learning rate for positive prediction errors; α-: average learning rate for negative prediction errors (Asymmetric Model); β_0_: softmax intercept (bias towards reject); β_1_: softmax slope (sensitivity to the difference in the value of rejecting versus the value of accepting an option). Data are expressed as mean and 95% confidence intervals (calculated as 1.96*standard error). Note: learning rates displayed here (α, α+ and α-) are untransformed parameters from the model fitting procedure; the function *0.5 + 0.5*erf(α/sqrt(2))* is subsequently applied to transform these to conventional learning rates within the range 0 to 1. \*\**p<0.01 comparing LOOCV scores between the two models, paired sample ttest*

| Model | LOOCV | $\alpha$ | $\alpha+$ | $\alpha-$ | $\beta_0$ | $\beta_1$ |
| --- | --- | --- | --- | --- | --- | --- |
| Symmetry Model | 1230.54 | -2.49<br>[95% CI:<br>$\pm 0.54$] | | | 1.80<br>[95% CI:<br>$\pm 0.66$] | 0.15<br>[95% CI:<br>$\pm 0.05$] |
| Asymmetry Model | 1076.02** | - | -2.17<br>[95%<br>CI:<br>$\pm 0.38$] | -2.38<br>[95%<br>CI:<br>$\pm 0.49$] | -0.15<br>[95% CI:<br>$\pm 1.79$] | 0.13<br>[95% CI:<br>$\pm 0.03$] |

Next, we looked to relate our fine-grained trialwise measures of participants physiological state to learning from positive (*α*^+^) and negative (*α*^−^) prediction errors. To do this, we tested two further models: (1) Asymmetry Phys *α*^+^; (2) Asymmetry Phys *α*^−^, separately for HR, PEP and HF HRV. Each model respectively allowed *α*^+^ or *α*^−^ to change trial as a function of participants trial to trial physiological state (see analysis branch 2 methods). Results are summarized in Table 2; in the case of HF HRV, we observed evidence that learning from both positive (*α*^+^) and negative (*α*^−^) prediction errors was modulated by the physiological state (both p-values below0.040; paired sample ttests comparing both Asymmetry HF HRV models versus Asymmetry model). The direction of the effect in both of these models was such that when HF HRV was increased, learning was reduced, i.e., *ρ* shifted upwards more slowly following reward, or downward more slowly following absence of reward. In the case of HR, we observed no evidence that learning from either prediction error was modulated by the physiological state (both p-values above 0.222). Finally, for PEP, we did not find evidence that learning from positive prediction errors (*α*^+^) was modulated by its state (*p* = 0.135), however we did find evidence that learning from negative prediction errors (*α*^−^) was modulated (*t*(19) = 2.18, *p* = 0.042; paired sample ttests comparing both Asymmetry PEP models versus Asymmetry model). The direction of the effect in this model was such that negative prediction errors that caused *ρ* to shift downwards (following the absence of a reward) changed beliefs to a greater extent when sympathetic drive was high (shorter PEP). In other words, participants were faster to learn that their environment was deteriorating when the sympathetic branch of the autonomic state was heightened.

**Table 2:**
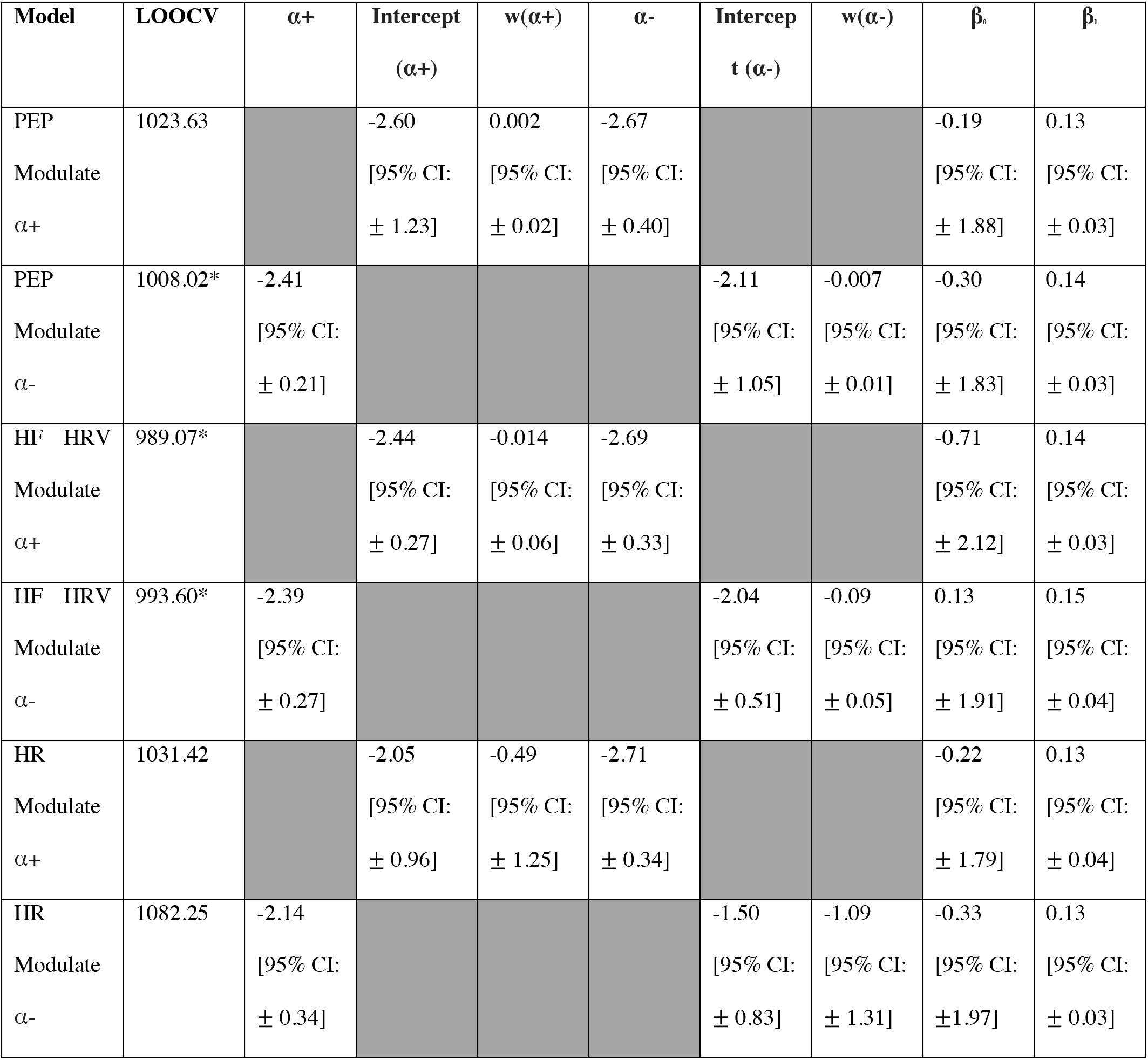
physiology-behaviour model fitting and parameters. model fitting and parameters for physiology modulated learning models. Each model allows either α+ or α- (untransformed) to be modulated on a trial by trial (t) basis according to one of the physiological readouts (PEP, HF HRV or HR) according to an intercept and a slope (e.g., α+(t) = intercept + w*PEP(t)). A transfer function [0.5 + *0.5*erf(α/sqrt(2))]* is then used to convert this to a conventional learning rate. The alternate learning rate is not modulated by physiology and is fit as a single free parameter. The table summarizes for each model its fitting performances and its average parameters: LOOCV: leave one out cross validation scores, summed over participants; α+: learning rate for positive prediction errors (untransformed); α-: average learning rate for negative prediction errors (untransformed); intercept: unstandardized intercept for regressing learning rate (α+ or α-) against physiology; w: unstandardized slope for regressing learning rate against physiology; β_0_: softmax intercept (bias towards reject); β_1_: softmax slope (sensitivity to the difference in the value of rejecting versus the value of accepting an option). Data are expressed as mean and 95% confidence intervals (calculated as 1.96*standard error). \**p<0.05 comparing LOOCV scores against scores for the basic Asymmetry Model which does not include physiology measures (see Table 1), paired sample ttest*

### Analysis branch 3 - Methods

Our analyses to this point reveal that perturbations in the reward rate are exclusively associated with changes in sympathetic tone, and that sympathetic activation increases learning symmetry between positive and negative information. We might accordingly expect sympathetic engagement to predict optimal performance in the prey selection task. Here we formalize optimal performance (*D – B Mid*) for each subject as the delta in the rate of mid-rank capture in the downturn environment, relative to the boom environment:

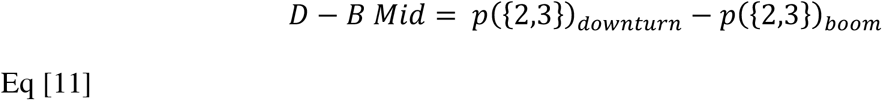

Where *p*(*X*)_*s*_ is the proportion of items in vector *X* captured in environment *S*. Higher positive *D – B Mid* values accordingly reflect more optimal prey selection in the downturn environment, which predicts better task performance. We also computed a similar downturn-boom delta value for each subject for each physiological environment:

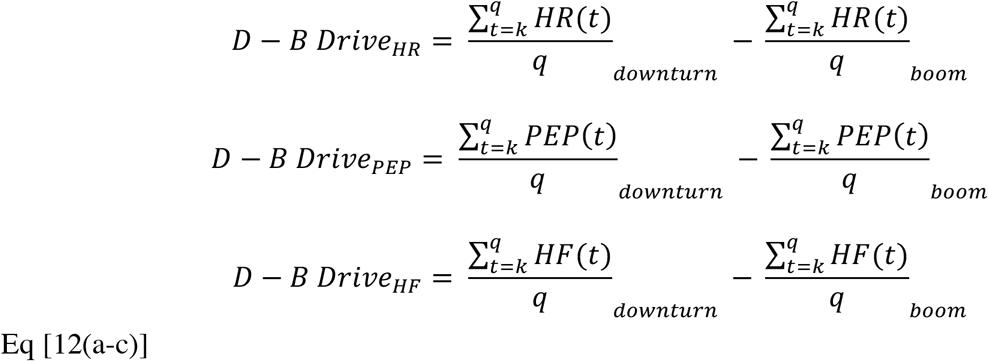

whereby each *D – B Drive* corresponds to the downturn-boom delta between the average of each trialwise physiological state from trial *k* to *q*. Higher positive values reflect higher drive (i.e. increased contractility, increased HF HRV power and increased HR) in the downturn environment. We probed the relationship between our assay of optimal performance and environment-wise physiology fluctuations with a linear model of each subject’s *D – B Mid* as a function of an intercept, and their *D – B Drive_HR_, D – B Drive_PEP_* and *D – B Drive_HF_* values. Further, we ran two iterations of this model, one from the trialwise measures taken from trials in the first half (i.e. 0-360s) of the time spent in each environment (where we assumed the majority of learning is required) and as a control, another model using trials in the second half (360-720s) of the time spent in each state. In other words, in Eq [12(a-c)], for the first model *k* was always 1 and *q* was the last trial for each subject that started before 360s. For the second model, *k* was the first trial that started after 360s, and *q* was the subject’s final trial. Positive coefficients associate the engagement of the relevant physiological state with optimized learning.

### Analysis branch 3 - Results

The linear model (see Fig. 3C) of *D – B Mid*, as a function of the *D – B Drive* value for each physiological state, estimated from trials in the first half (early) of each reward state returned a significant positive coefficient for *D – B Drive_PEP_* (*β* = 0.035, *s.e*. = 0.016, *p* = 0.046), with neither coefficient for *D – B Drive_HR_* nor *D – B Drive_HF_* reaching statistical significance (both p-values > 0.569). The model of *D – B Mid* using physiological *D – B Drive* values estimated from trials in the second half (late) of reward states did not return any significant coefficients (all p-values > 0.134). These final models suggest that the sympathetic engagement during crucial learning periods of a low reward environment predicts optimal behavioral adjustment.

## Discussion

Appraising sequential offers of reward relative to an unknown future opportunity and a time cost requires an optimization policy that draws on a belief about the richness characteristics of the current environment. Across a range of experiments, including reinforcement-learning tasks, belief updating paradigms and prey selection, information integration shows a positive bias (Garrett and Sharot, 2014, 2017; Garrett et al., 2014, 2018; Kuzmanovic et al., 2015, 2016; Eil and Rao, 2011; Korn et al., 2012; Kuzmanovic and Rigoux, 2017; Lefebvre et al., 2017; Garrett and Daw, 2019). That is, the rate at which humans update their belief about a probability, association or reward rate is more sluggish if the new information carries a negative or aversive valence or telegraphs a deterioration of the current belief. In our prey selection task, subjects updated their belief about the rate of harvested reward with a similar bias toward reward over delay information. However, using simultaneous continuously recorded cardiac autonomic physiology measures, we reveal a uniquely adaptive role for the sympathetic branch in this situation.

In our choice models (analysis 1), activation in both branches of the autonomic state blunted the positive association between value and choice, whether or not the immediate context was factored into the value of the offer, i.e., whether cost reflected the objective capture-time of an invader and reward reflected the objective harvested fuel, or whether these value dimensions were collapsed into a single variable that compared reward to the opportunity cost of capture in the current reward context. However, we observed contextual derivatives, i.e., changes in the rate of harvested reward, having a unique association with sympathetic over parasympathetic state derivatives. Specifically, increases in contractility (shorter PEP) scaled with decreases in the average rate of reward harvested per second. No relationship emerged between contractility changes and absolute fluctuations in the environment’s richness, dissociating sympathetic drive from a more global richness-monitoring role that tracks outright contextual changes regardless of direction or valence. In our learning models (analysis 2), drive in the sympathetic system uniquely increased the rate of learning, and specifically when reward rate estimates were updated via negative prediction error. In contrast, parasympathetic drive aligned with decreased learning rates from both positive and negative prediction error. Finally, in analysis 3, we revealed that the unique deterioration specific relationship between sympathetic drive and learning was not exclusively a phenomenological response to a worsening environment, by observing a positive relationship between activation of the sympathetic state during crucial periods of the task – i.e., early in the downturn environment - and deployment of optimal behavioral policy – i.e., increased capture of mid rank invaders.

The only other study (Lenow et al., 2017) to probe the stress system in a human foraging task demonstrated an opposing relationship whereby stress increased overharvesting. Overharvesting occurs when an animal or human exploits proximal known resources beyond a threshold determining that better yields would be obtained by switching location. Such maladaptive perseveration indicates a biased low estimation of environmental quality. That this tendency is exacerbated by stress is interestingly both consistent and at odds with our finding that sympathetic stress adaptively increases the rate of negative information integration. Consistent, in so far as stress drives a pessimistic learning style in both cases, but at odds in terms of adaptivity. It could be the case that stress simply plays a general role in driving more pessimistic foraging behavior and whether or not this proves adaptive is an arbitrary consequence of task design. However it’s important to also note that Lenow et al. (2017) assayed the hypothalamic-pituitary-adrenal axis of the stress response (HPA) via cortisol. Some studies show alignment between the sympathetic system and the HPA response to stressors (e.g., Bosch et al., 2009), others demonstrate task selectivity, particularly where subjects feel threat or a loss of control (see Dickerson and Kemeny (2004) for review) and others still argue a sequential framework that sees HPA respond once a threshold level of activation has been reached by the sympathetic system (Cacioppo et al., 1995; Bosch et al., 2009).

The unique reactivity of the sympathetic branch of the autonomic system for learning that the rate of reward is deteriorating is consistent with division specific neural control of autonomic function. Meta-analysis of human neuroimaging data demonstrate that divergent brain networks regulate the sympathetic and parasympathetic branch, with control of the former including prefrontal and insular cortices, in addition to multiple areas within the medial wall of the mid and anterior cingulate (ACC; Beissner et al., 2013; Dum et al., 2016). Further, animal tracer evidence (Dum et al., 2016) reveals direct synaptic inputs into the adrenal medulla - a key sympathetic site for catecholaminergic release - from both pregenual and subgenual portions of ACC. The specific mobilization of the sympathetic branch when the reward rate is deteriorating may offer a support mechanism for involvement of the ACC, a key node in a network associated with value-based decisions, specifically decisions that involve costs or negative affect (Walton et al., 2009, Shackman et al., 2011, Amemori and Graybiel, 2012). ACC has also been implicated amongst other prefrontal sites in tracking reward history (Seo et al., 2007; Bernacchia et al., 2011). This would align with previous neurophysiological work in human learning, which implicates heightened stress states – assayed via electrodermal activity - with closer trialwise tracking of adjustments in information integration (Li et al., 2011), with selective stress-driven sensitivity increases to negative prediction errors, eradicating positive learning biases that emerge under calm conditions when stress levels are normal (Garrett et al., 2018).

A curious finding emerged in our initial choice models, where drive in both sympathetic and parasympathetic branches was associated with low value capture; curious given the antagonistic manner in which the two branches of the autonomic state typically respond, e.g., a stressor will induce increased contractility of the heart and decreased vagal tone. This parallel activation could therefore reflect the two autonomic branches serving different functions in the prey selection task via different time constants. As predicted by MVT, and as evidenced by increases in mid-rank invader captures, the downturn reward state increased the subjective value of lower value items. Thus, having learned of a deteriorated environmental richness (associated with increased sympathetic activation), subjects now require a change in behavior policy. In other words, subjects must overwrite the prepotent policy of avoiding mid-rank invaders, and switch to a policy of capturing them. Parasympathetic drive may therefore signal increased executive control requirements. Neurovisceral integration models of cognitive control (Thayer et al., 2009) would support this interpretation. Under this model, prefrontal-subcortical inhibitory circuits that govern the control of thoughts and goal-directed behavior provide inhibitory input to the heart via the vagus nerve (Benarroch, 1993; Ellis and Thayer, 2010). Later studies supported this model by demonstrating participants with higher resting state HF HRV perform better on task switching (Colzato et al., 2018), supporting the idea that vagal tone is important for adaptive responses to environmental demands (Thayer and Lane, 2000). Human meta-analytic imaging data also support a vagal role in supporting behavioral adaptation; sympathetic neural sites are more associated with paradigms involving cognitive stress, while parasympathetic neural sites contribute where tasks involve somatosensory-motor performance (Beissner et al., 2013). While no evidence exists in humans for such parallel autonomic drive, examples exist in the animal literature, such as parallel sympathetic and vagal outflow during the freeze response to conditioned fearful stimuli (Carrive, 2006).

However, it’s important to note that an alternative to this parallel processing framework is that the shorter PEP associated with low value capture was not driven by sympathetic drive, but was instead a downstream consequence of a dominant parasympathetic engagement. This interpretation is supported by the additional association we observed between decreased heart-rate and low value capture. Heart-rate can link the concurrent sympathetic and parasympathetic associations via the Frank-Starling mechanism, whereby a beat to beat deceleration in heart rate, which increases the preload, i.e., ventricular filling, causes increased contractility (shorter PEP) for reasons not mediated by the sympathetic system (Sherwood et al., 1990; Kuipers et al., 2017). In other words, capture of low value may be predominantly associated with increased vagal tone, which transiently slows the heart-rate, which in turn increases PEP via increased preload. Such an autonomic cascade would align with the directions we observed in the coefficients relating the physiology variables and value in our choice models.

Ultimately, we can only speculate at this point with regard to the precise roles of the parasympathetic and sympathetic branch in making low value choices, and an aim for future research should be tasks that manipulate both learning and executive demands while recording simultaneous measures of both branches of the autonomic state. It’s nonetheless important to clarify that the Frank-Starling effect does not apply to the primary association revealed by our learning models, models of reward rate derivatives, and model of choice optimization, all of which did not show concurrent influence of heart rate with other autonomic responses, and all of which point to the sympathetic system being uniquely involved in tracking deteriorations in reward rate, and that sympathetic drive during crucial learning windows is uniquely associated with optimal behavioral adaptation in the prey selection task.

Cannon originally proposed that the body readies itself for a ‘fight or flight’ via secretions of the adrenal medulla, initiated by sympathetic neural projections from the thoracic spine. Our findings offer a potential parcellation of the modulatory role of the autonomic system in context specific decisions - sympathetic support of learning environmental deterioration, and, possibly, parasympathetic support of behavior modification.

## Acknowledgements

The research was supported by award #W911NF-16-1-0474 from the Army Research Office and by the Institute for Collaborative Biotechnologies under Cooperative Agreement W911NF-19-2-0026 with the Army Research Office. Authors are indebted to Tom Bullock and Alexandra Stump for technical assistance and data collection.

